## Supplementary figures and images for "Temporal pattern and synergy influence activity of ERK signaling pathways during L-LTP induction"

### Figure 2 -supplemental figure 1

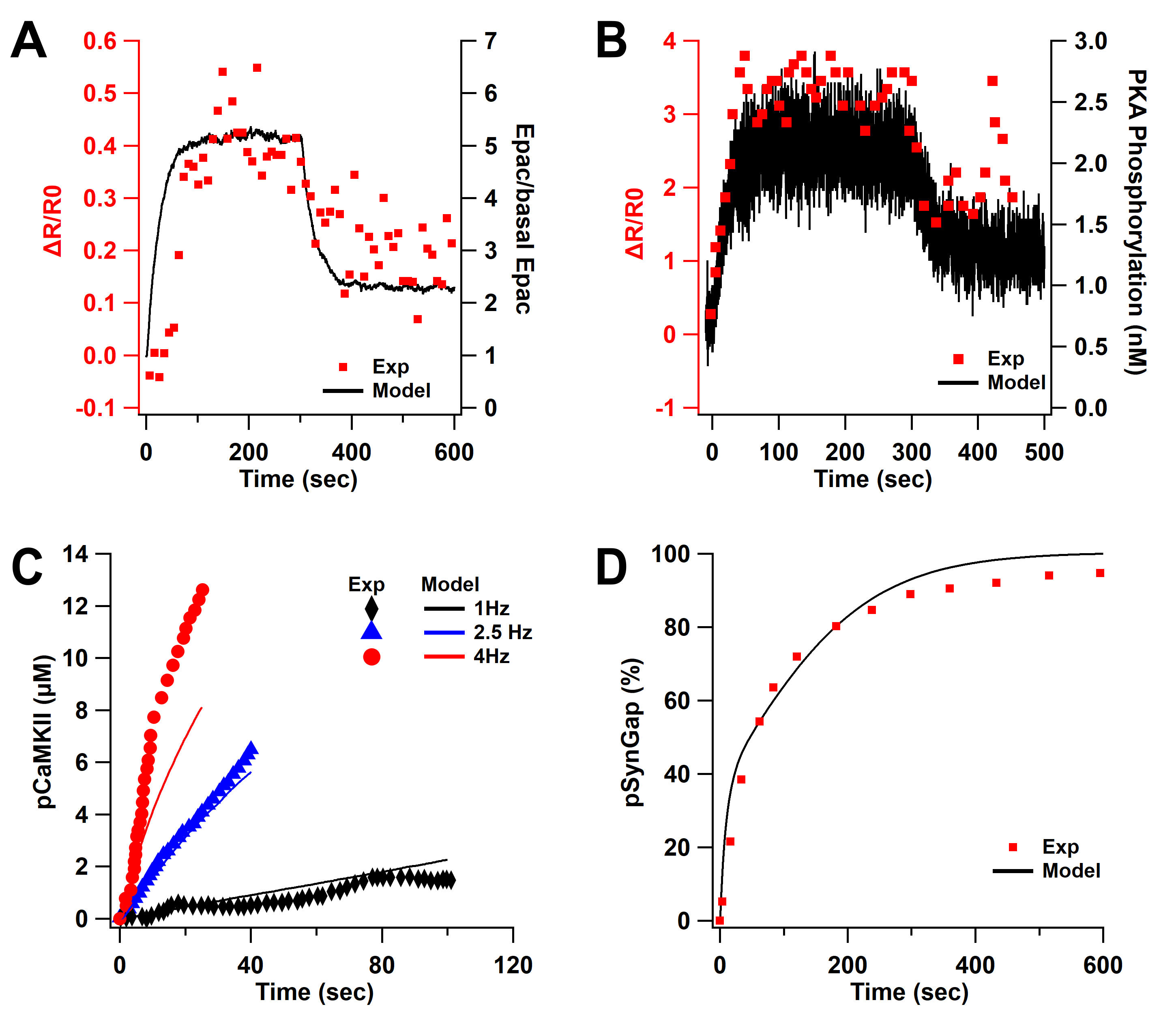

### Figure 3 -supplemental figure 1

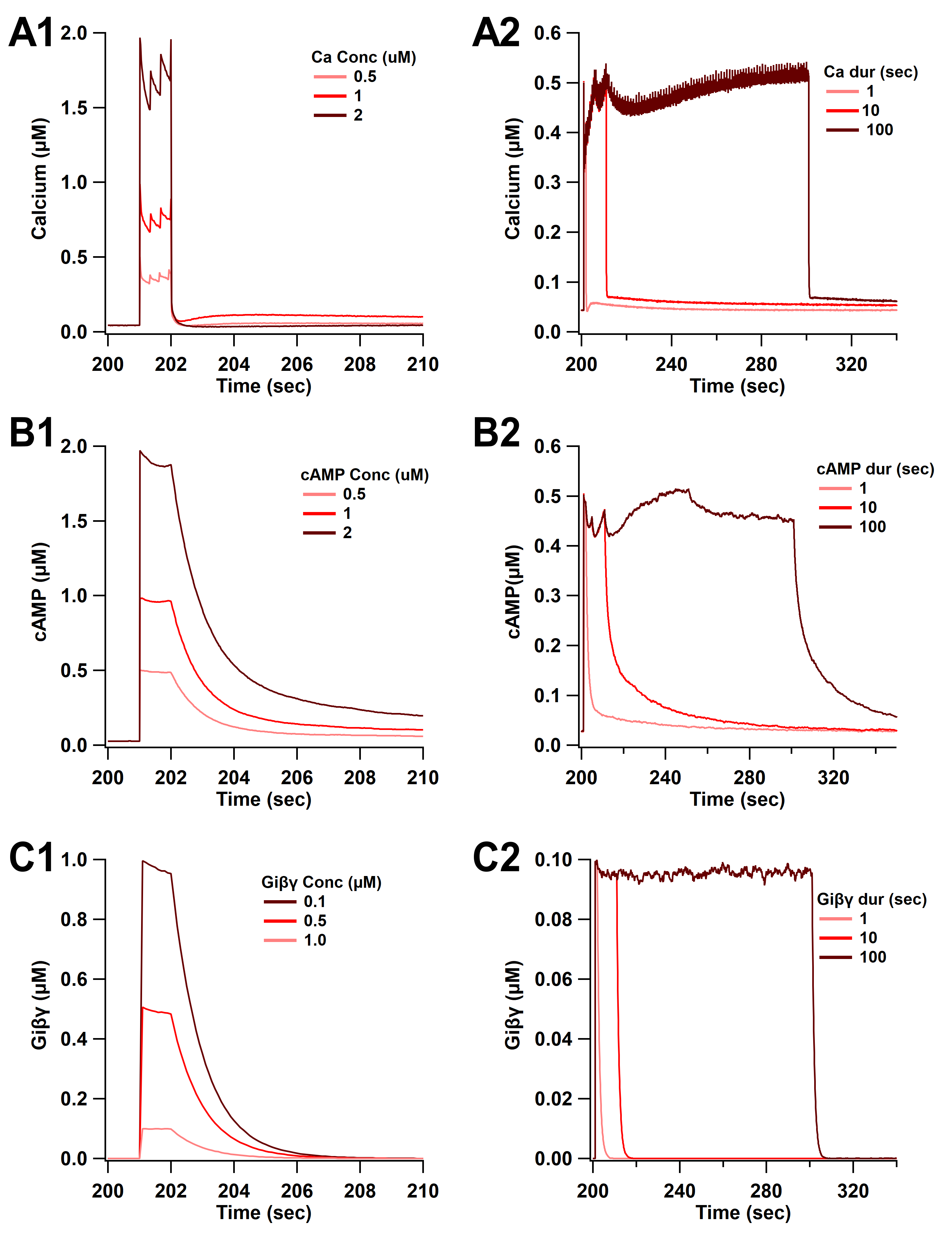

### Figure 6 -supplemental figure 1

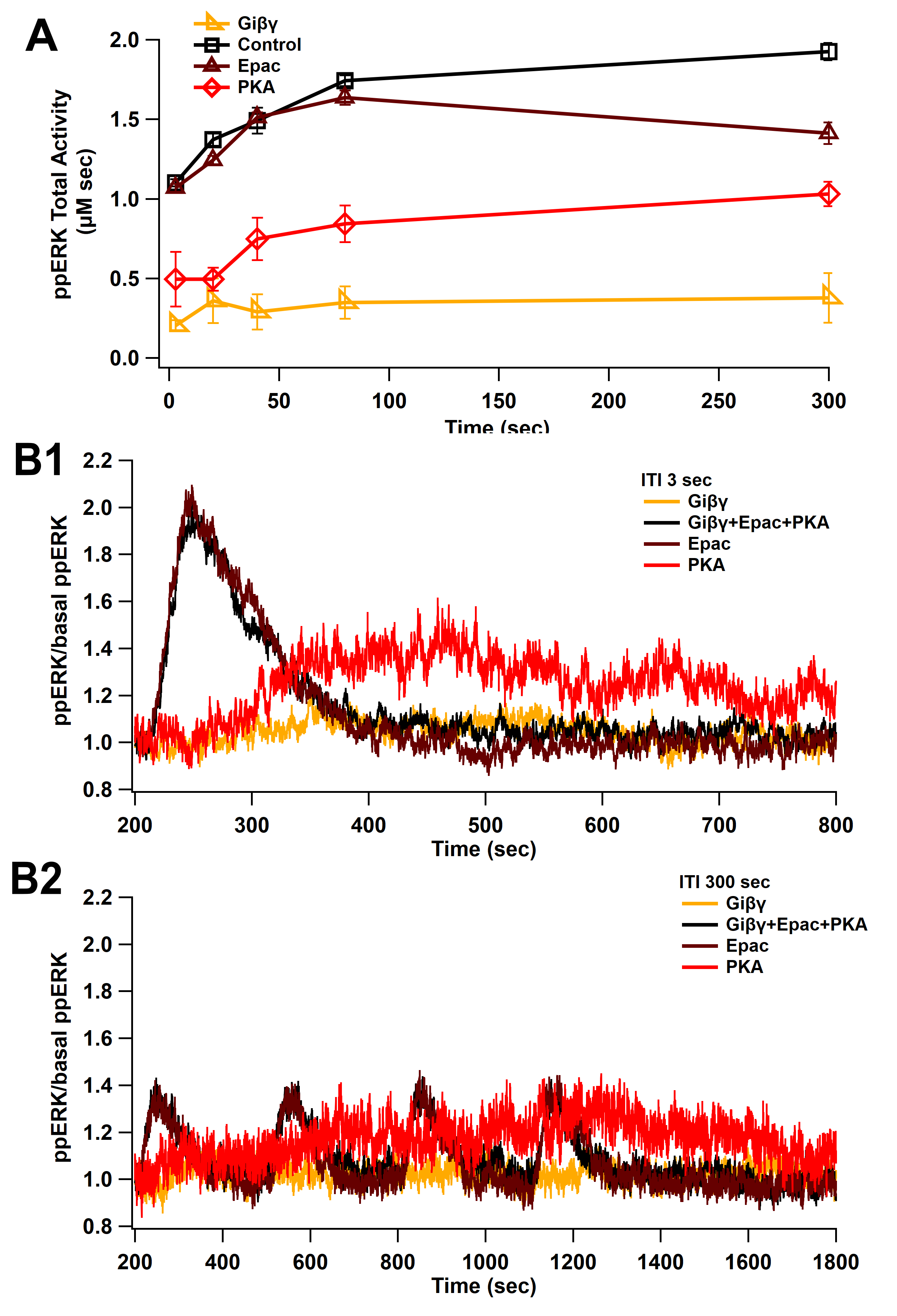

### Figure 8 -supplemental figure 1

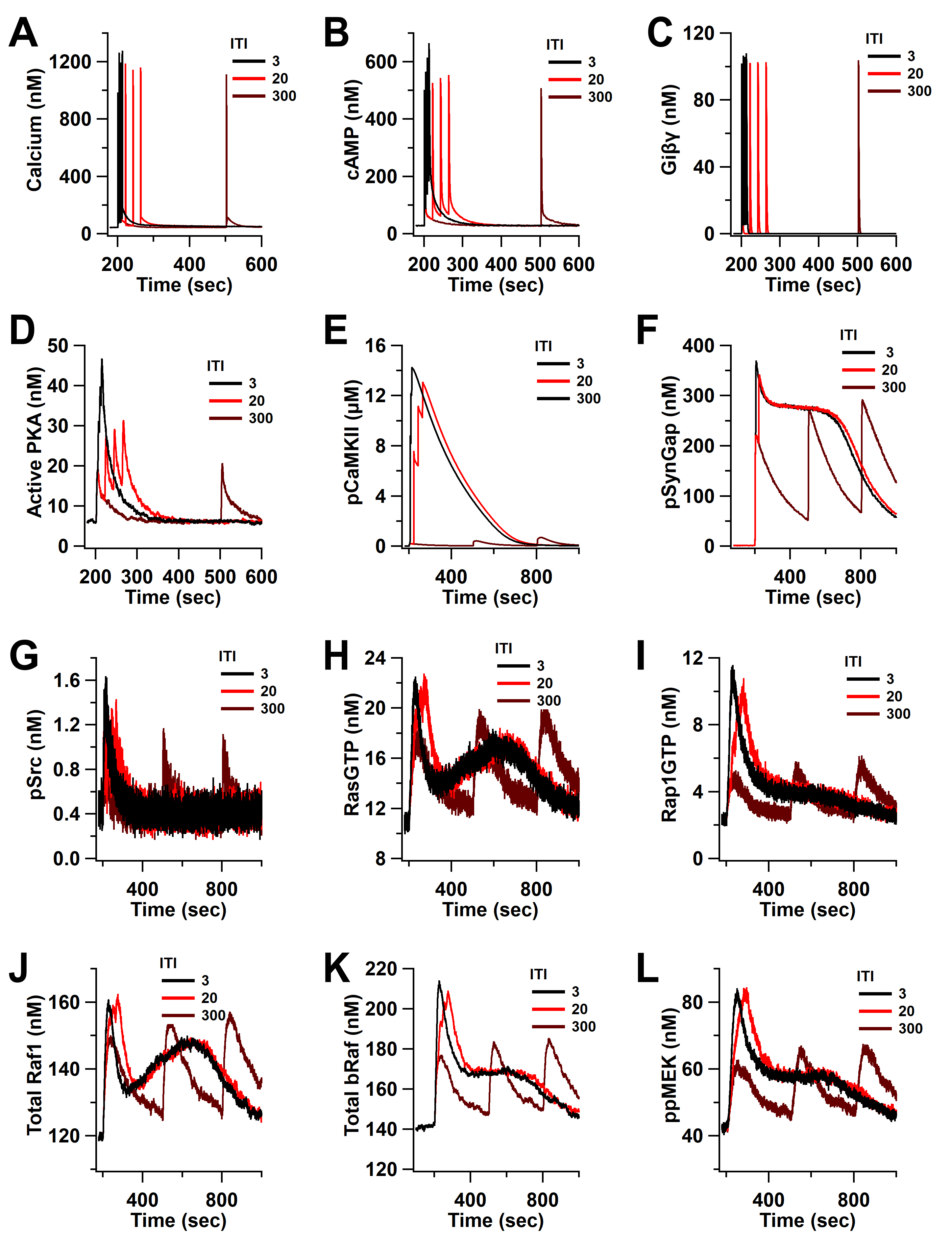

### Figure 8 -supplemental figure 2

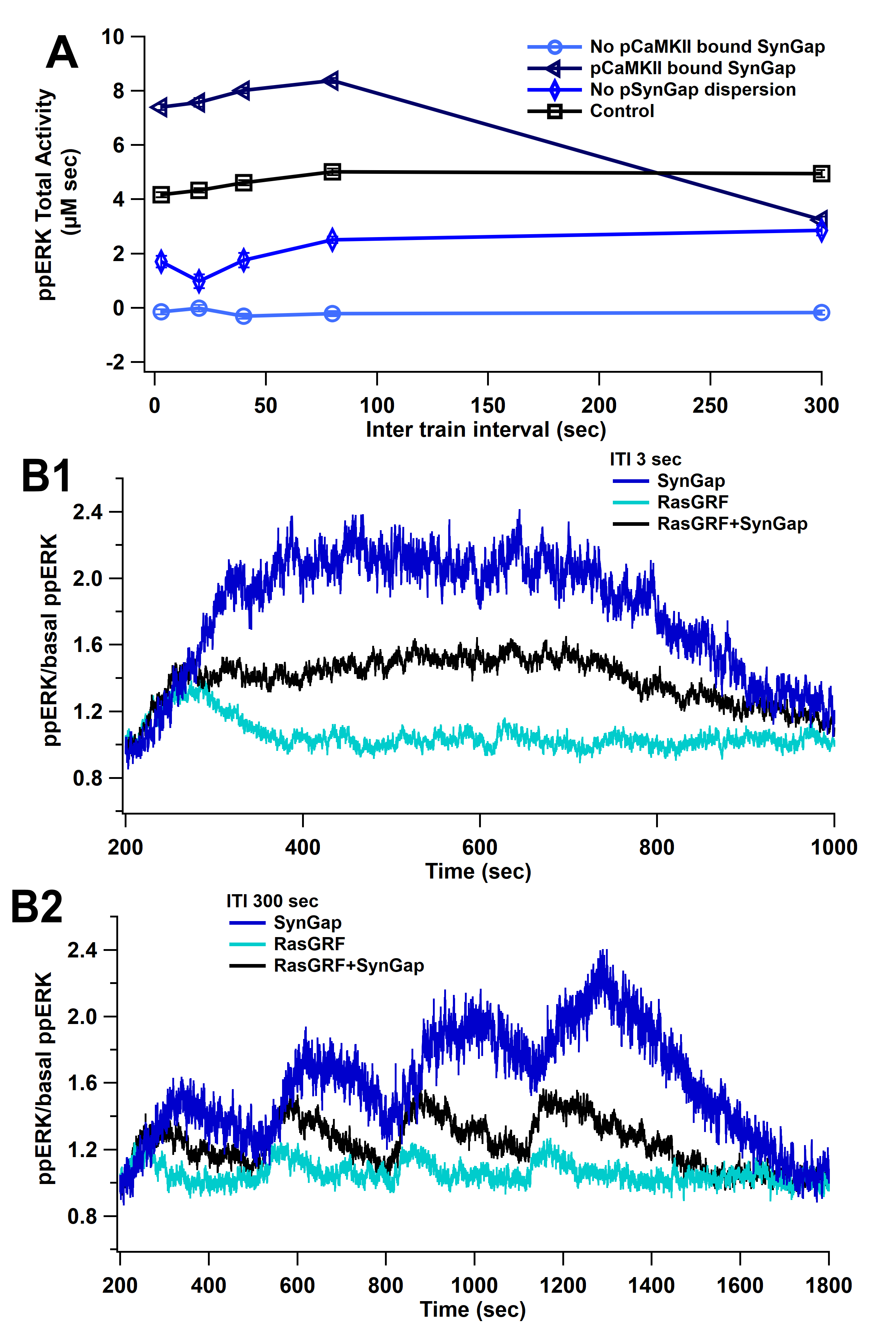

### Figure 8 -supplemental figure 2

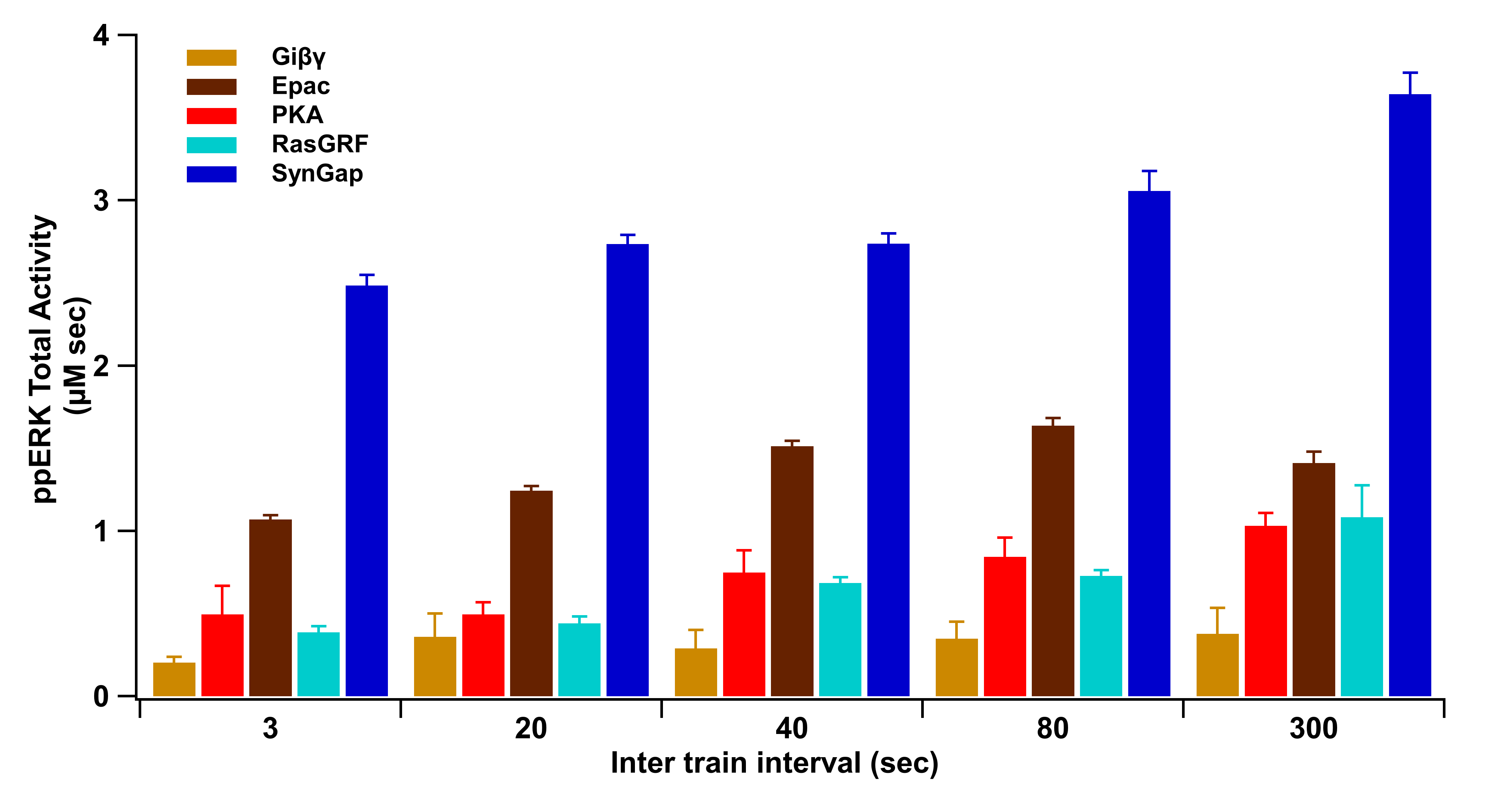

### Figure 8 -supplemental figure 3

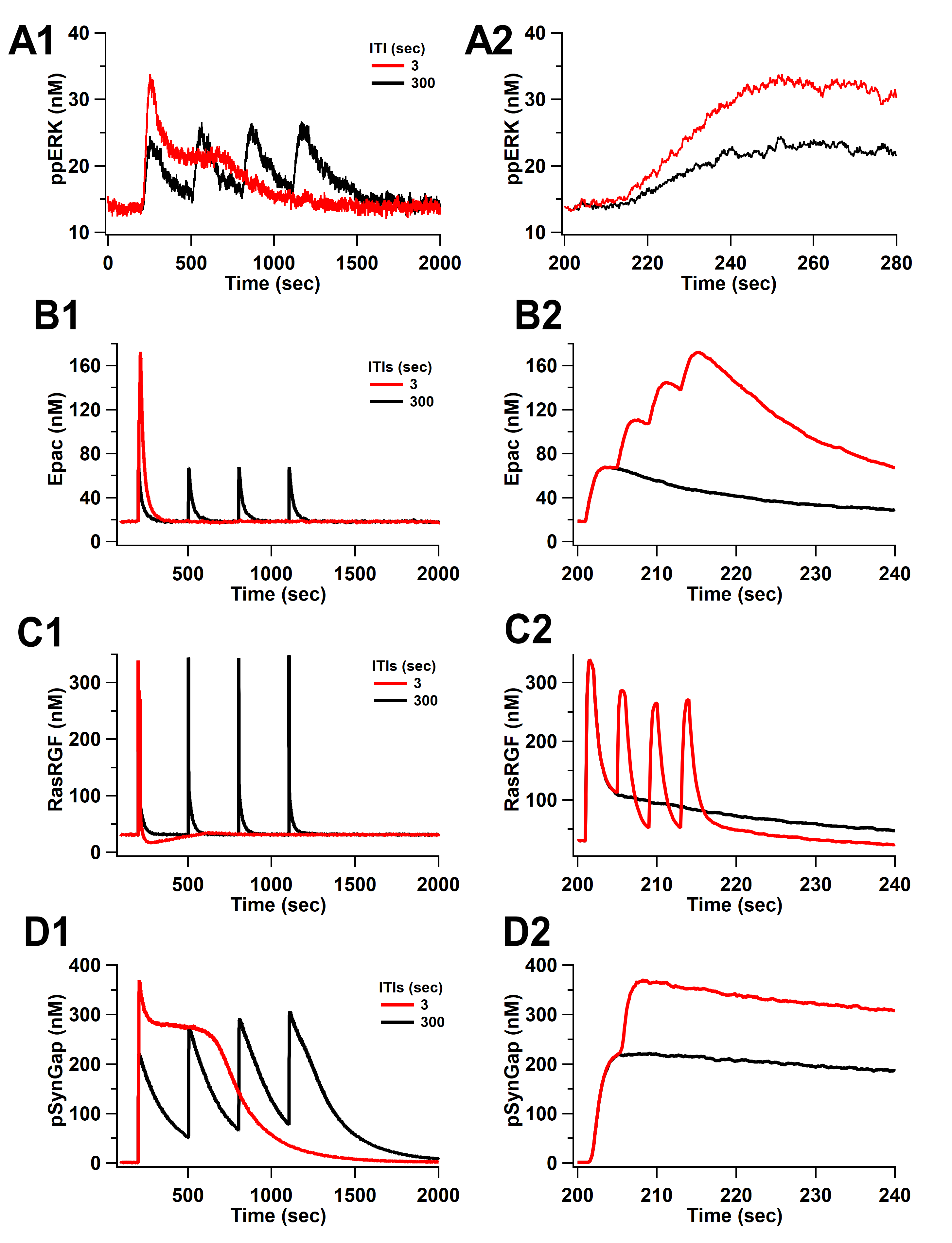

### Figure 8 -supplemental figure 4

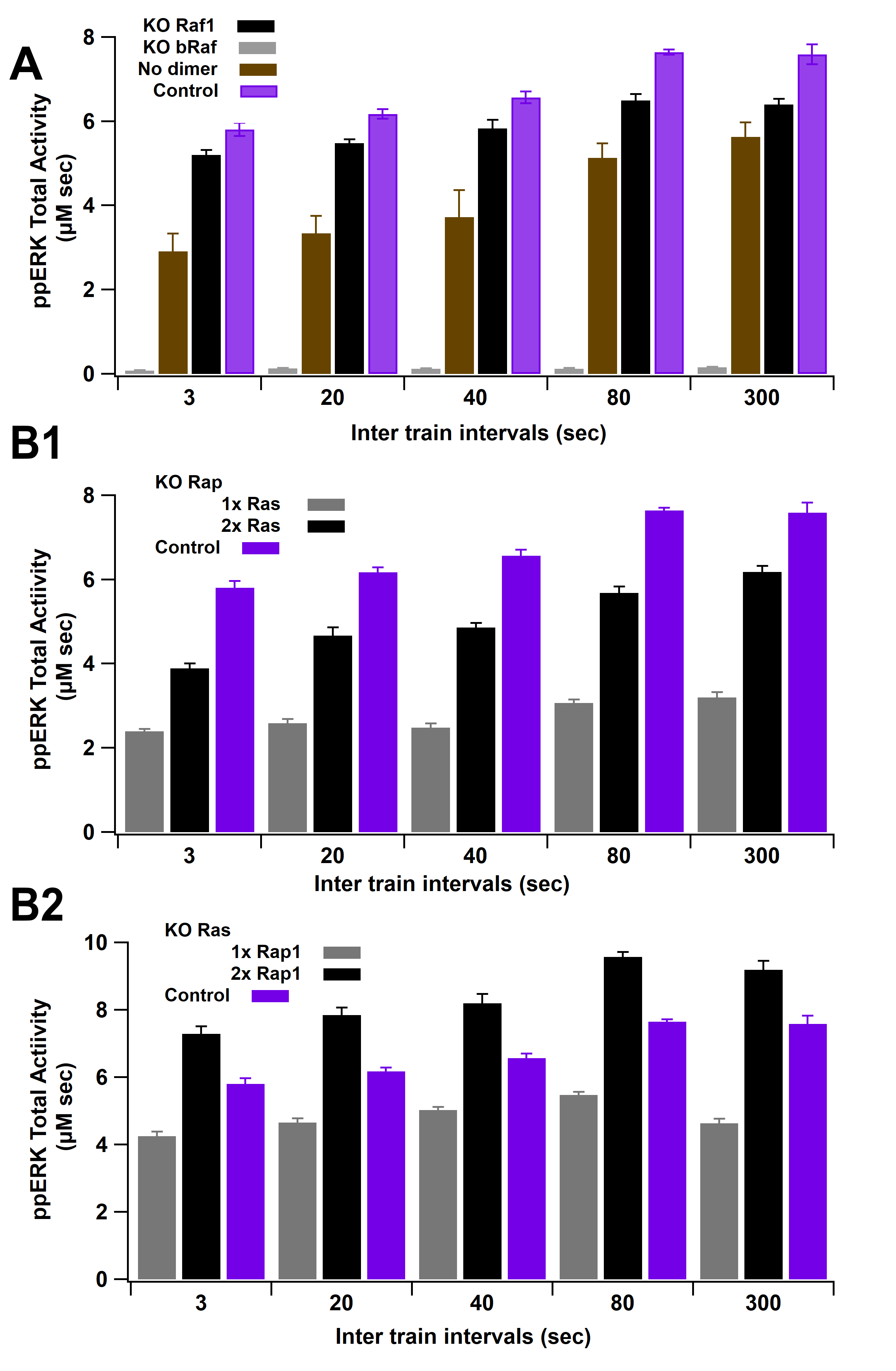

### Figure 9 -supplemental figure 1

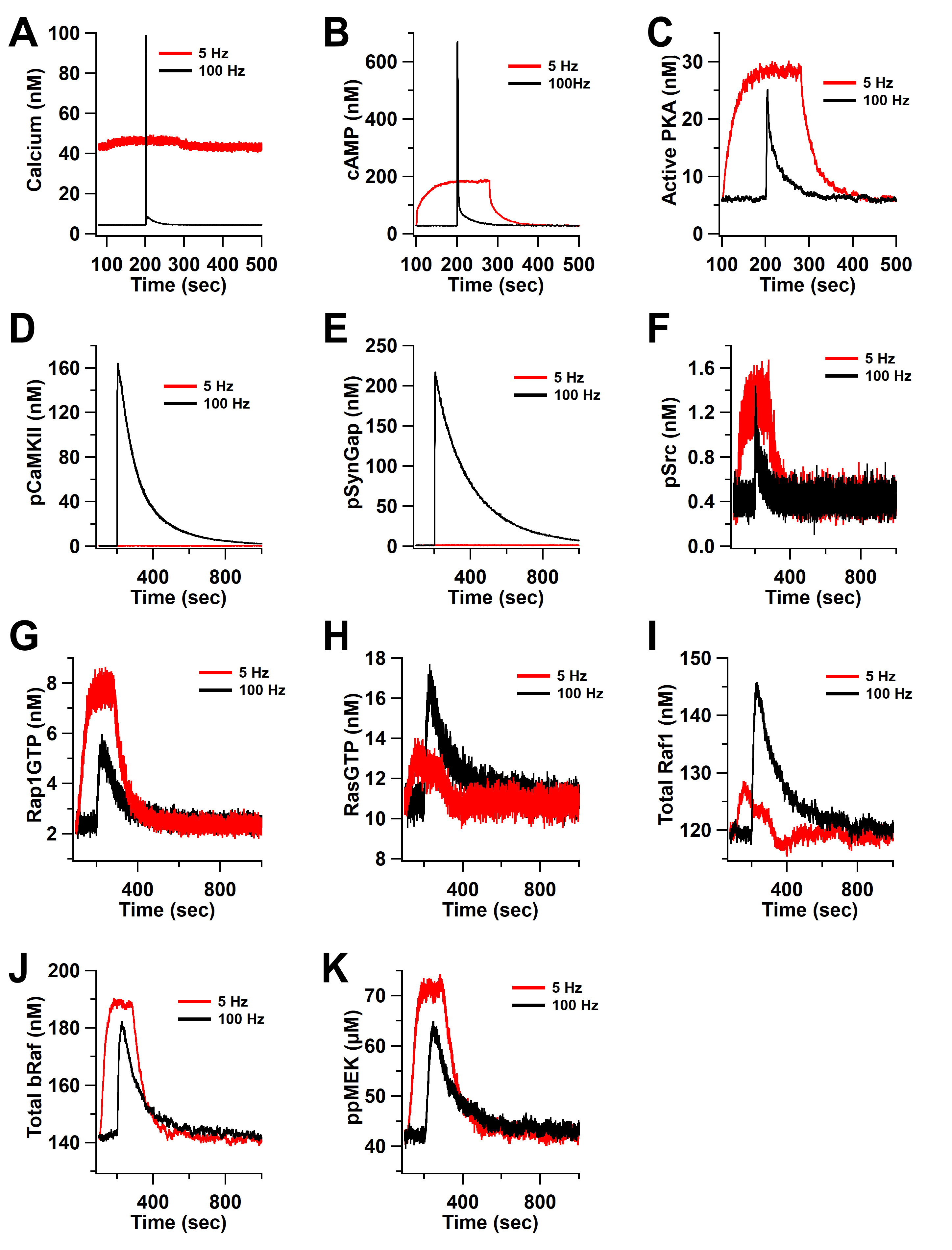
