## Supplementary material for "Temporal pattern and synergy influence activity of ERK signaling pathways during L-LTP induction": Figure 7 -supplemental figure 1

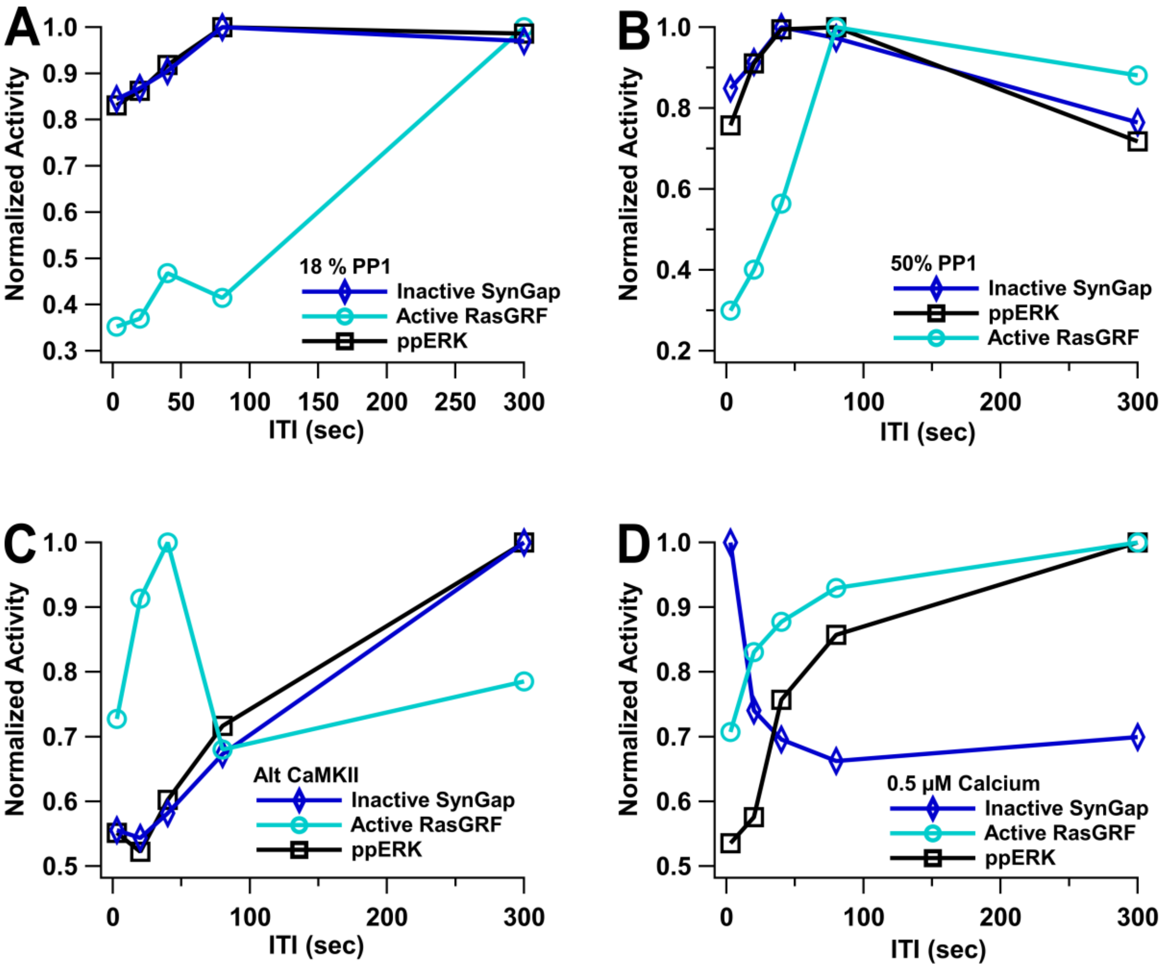


**E**

| | | Correlation | |
|---|---|---|---|
| | | Inactive SynGap | Active RasGRF |
| 18% PP1 | R | 0.9907 | 0.5832 |
| | pValue | **0.0005** | 0.1510 |
| 50% PP1 | R | 0.9693 | 0.1852 |
| | pValue | **0.0032** | 0.3827 |
| alt CaMKII | R | 0.9917 | -0.3188 |
| | pValue | **0.0004** | 0.6994 |
| 0.5 µM calcium | R | -0.6971 | 0.9450 |
| | pValue | 0.9046 | **0.0077** |
