## Supplemental Table 1 for "Temporal pattern and synergy influence activity of ERK signaling pathways during L-LTP induction"

**Figure 1 –** **Source Data 1:** Reactions and rates constant involved in signaling pathway leading to Ras and Rap1 activation

| Reaction equation | K_f_ (nM^-1^ Sec^-1^) | | K_b_ (Sec^-1^) | K_cat_ (Sec^-1^) | Reference |
| --- | --- | --- | --- | --- | --- |
| *Gby + Src* $\boldsymbol{\leftrightarrow}$ *Src_Gby* | | 2.00E-04 | 1.00E-01 |  | Estimated |
| *Grb2 + Sos* $\boldsymbol{\leftrightarrow}$ *Grb2_Sos* | | 2.50E-04 | 1.68E-02 |  | Chook et al., 1996; Jain and Bhalla, 2014 |
| *Src_Gby +Shc* $\boldsymbol{\leftrightarrow}$ *pShc +Src_Gbg* | | 8.00E-03 | 1.28E-01 | 3.20E-02 | Estimated |
| *pShc + Src_Grb2_Sos* $\boldsymbol{\leftrightarrow}$ *pShc_Grb2_Sos* | | 5.00E+00 | 1.00E-01 |  | Bhalla and Iyengar, 1999; Jain and Bhalla, 2014; Sasagawa et al., 2005 |
| *pShc_Grb2_Sos + RasGDP* $\boldsymbol{\leftrightarrow}$ *pShc_Grb2_Sos + RasGTP* | | 1.98E-02 | 8.00E-01 | 2.00E-01 | Bhalla and Iyengar, 1999; Jain and Bhalla, 2014; Sasagawa et al., 2005 |
| *pShc* $\boldsymbol{\leftrightarrow}$ *Shc* | | 2.00E-01 | 0.00E+00 |  | Jain and Bhalla, 2014 |
| *ppERK +pShc_Grb2_Sos*$\boldsymbol{\leftrightarrow}$*ppERK +pShc_Grb2 + pSos* | | 1.95E-02 | 4.00E+01 | 1.00E+01 | Jain and Bhalla, 2014; Sasagawa et al., 2005 |
| *pSos*$\boldsymbol{\leftrightarrow S}$*os* | | 1.00E+00 |  |  | Jain and Bhalla, 2014 |
| *CaMCa4 + Ras-GRF2* $\boldsymbol{\leftrightarrow}$ *Ras-GRF2_CaMCa4* | | 6.00E-02 | 9.24E-01 |  | Jin et al., 2014 |
| *Ras-GRF2_CaMCa4 + RasGDP* $\boldsymbol{\leftrightarrow}$ *Ras-GRF2_CaMCa4 + RasGTP* | | 1.98E-02 | 8.00E-01 | 2.00E-01 | Farnsworth et al., 1995; Jin et al., 2014, |
| *PKAc + Src* $\boldsymbol{\leftrightarrow}$ *pSrc + PKAc* | | 2.04E+00 | 8.00E+01 | 2.00E+01 | Jain and Bhalla, 2014 |
| *pSrc + Cbl* $\boldsymbol{\leftrightarrow}$ *pCbl + Src* | | 4.00E-01 | 1.60E+02 | 4.00E+01 | Jain and Bhalla, 2014 |
| *Crk_C3G + pCbl* $\boldsymbol{\leftrightarrow}$*Crk_C3G_pCbl* | | 1.00E-03 | 2.00E-01 |  | Jain and Bhalla, 2014; Knudsen et al., 1994; Sasagawa et al., 2005 |
| *pCbl* $\boldsymbol{\leftrightarrow}$*Cbl* | | 1.00E+02 |  |  | Jain and Bhalla, 2014 |
| *pSrc* $\boldsymbol{\leftrightarrow}$*Src* | | 1.00E+02 |  |  | Jain and Bhalla, 2014 |
| *Crk_C3G_pCbl + Rap1GDP* $\boldsymbol{\leftrightarrow}$*Crk_C3G_pCbl + Rap1GTP* | | 1.01E-02 | 8.00E-02 | 2.00E-02 | Gotoh et al., 1995; Jain and Bhalla, 2014; Sasagawa et al., 2005 |
| *Epac + cAMP* $\boldsymbol{\leftrightarrow}$ *Epac_cAMP* | | 1.21E-04 | 1.45E-01 |  | de Rooij et al., 2000; Luczak et al., 2017 |
| *Epac_cAMP + Rap1GDP* $\boldsymbol{\leftrightarrow}$ *Epac_cAMP + Rap1GTP* | | 1.20E-03 | 9.60E-01 | 2.40E-01 | Estimated |
| *Rap1GTP+Rap1Gap* $\boldsymbol{\leftrightarrow}$ *Rap1GDP* | | 2.00E-01 | 2.00E+02 |  | Daumke et al., 2004; Jain and Bhalla, 2014; Sasagawa et al., 2005 |
| *RasGTP+ RasGap* $\boldsymbol{\leftrightarrow}$ *RasGDP* | | 4.95E-02 | 4.00E+01 |  | Bhalla and Iyengar, 1999; Jain and Bhalla, 2014 |
