## Supplemental Table 2 for "Temporal pattern and synergy influence activity of ERK signaling pathways during L-LTP induction"

**Figure 1 –** **Source Data 2:** Reaction and rates constant involved in signaling pathway of SynGap. Rates were optimized from Oh et al., 2004 and Walkup IV et al., 2015 (Miningou and Blackwell, 2018).

| Reaction equation | K_f_ (nM^-1^ Sec^-1^) | K_b_ (Sec^-1^) | K_cat_ (Sec^-1^) | Reference |
| --- | --- | --- | --- | --- |
| *CKpCaMCa4 + SynGAP* $\boldsymbol{\leftrightarrow}$ *pSynGAP + CKpCaMCa4* | 3.07E-01 | 5.87E+01 | 6.48E-01 | Miningou and Blackwell, 2018 |
| *Ras1GTP + SynGAP* $\boldsymbol{\leftrightarrow}$ *Ras1GDP + SynGAP* | 8.20E-04 | 8.20E-01 | 2.05E-01 | Miningou and Blackwell, 2018 |
| *Rap1GTP + SynGAP* $\boldsymbol{\leftrightarrow}$ *Rap1GDP + SynGAP* | 3.20E-03 | 3.20E+00 | 8.00E-01 | Miningou and Blackwell, 2018 |
| *RasGTP + pSynGAP* $\boldsymbol{\leftrightarrow}$ *RasGDP + pSynGAP* | 1.12E-03 | 1.11E+00 | 2.79E-01 | Miningou and Blackwell, 2018 |
| *Rap1GTP + pSynGAP* $\boldsymbol{\leftrightarrow}$ *Rap1GDP + pSynGAP* | 6.14E-03 | 6.14E+00 | 1.53E+00 | Miningou and Blackwell, 2018 |
| *pSynGAP* $\boldsymbol{\leftrightarrow}$ *SynGAP* | 1.00E+00 | 0.00E+00 |  | Estimated |
| *pSynGAP* $\boldsymbol{\leftrightarrow S}$*ynGap_dendrite* | 2.50E-02 | 2.50E-02 |  | Araki et al., 2015 |
