## Supplemental Table 3 for "Temporal pattern and synergy influence activity of ERK signaling pathways during L-LTP induction"

**Figure 1 – Source Data 3:** Reactions and rates constant involved in core ERK signaling pathway

| Reaction equation | K_f_ (nM^-1^ Sec^-1^) | K_b_ (Sec^-1^) | K_cat_ (Sec^-1^) | Reference |
| --- | --- | --- | --- | --- |
| *Rap1GTP + B-Raf* $\boldsymbol{\leftrightarrow}$ *B-Raf_Rap1GTP* | 6.00E-02 | 1.00E+00 |  | Jain and Bhalla, 2014; Sasagawa et al., 2005 |
| *Rap1GTP + Raf-1* $\boldsymbol{\leftrightarrow}$ *RapGTP_Raf-1 + RasGTP* | 6.00E-03 | 5.00E-01 |  | Jain and Bhalla, 2014; Sasagawa et al., 2005 |
| *B-Raf_Rap1GTP + MEK* $\boldsymbol{\leftrightarrow}$ *pMEK + B-Raf_Rap1GTP* | 9.38E-03 | 1.20E+00 | 3.00E-01 | ain and Bhalla, 2014; Sasagawa et al., 2005 |
| *B-Raf_Rap1GTP + pMEK* $\boldsymbol{\leftrightarrow}$ *ppMEK + B-Raf_Rap1GTP* | 9.38E-03 | 1.20E+00 | 3.00E-01 | Jain and Bhalla, 2014; Sasagawa et al., 2005 |
| *RasGTP + Raf1* $\boldsymbol{\leftrightarrow}$ *Raf1_ RasGTP* | 6.00E-02 | 1.00E+00 |  | Block et al., 1996; Force et al., 1994; Jain and Bhalla, 2014 |
| *2Raf1_ RasGTP* $\boldsymbol{\leftrightarrow}$*dRaf1_ RasGTP* | 1.00E-02 | 5.00E-01 |  | Estimated |
| *RasGTP + bRaf* $\boldsymbol{\leftrightarrow}$ *bRaf_RasGTP* | 6.00E-03 | 5.00E-01 |  | Jain and Bhalla, 2014; Yamamori et al., 1995 |
| *dRaf1_RasGTP + MEK* $\boldsymbol{\leftrightarrow}$ *pMEK + dRaf1_RasGTP* | 1.89E-02 | 2.40E+00 | 6.00E-01 | Estimated |
| *dRaf1_RasGTP + pMEK* $\boldsymbol{\leftrightarrow}$ *ppMEK + dRaf1_RasGTP* | 1.89E-02 | 2.40E+00 | 6.00E-01 | Estimated |
| *B-Raf _RasGTP + MEK* $\boldsymbol{\leftrightarrow}$ *pMEK + B-Raf _RasGTP* | 6.29E-03 | 8.00E-01 | 2.00E-01 | Jain and Bhalla, 2014; VanScyoc et al., 2008 |
| *B-Raf _RasGTP + pMEK* $\boldsymbol{\leftrightarrow}$ *ppMEK + B-Raf _RasGTP* | 6.29E-03 | 8.00E-01 | 2.00E-01 | Jain and Bhalla, 2014; VanScyoc et al., 2008 |
| *ppMEK + PP2A* $\boldsymbol{\leftrightarrow}$ *pMEK + PP2A* | 1.92E-03 | 2.40E+01 | 6.00E+00 | Jain and Bhalla, 2014; Takai and Mieskes, 1991 |
| *ppMEK + PP2A* $\boldsymbol{\leftrightarrow}$ *MEK + PP2A* | 1.92E-03 | 2.40E+01 | 6.00E+00 | Jain and Bhalla, 2014; Takai and Mieskes, 1991 |
| *ppMEK + ERK* $\boldsymbol{\leftrightarrow}$ *pERK + ppMEK* | 3.24E-02 | 1.20E+00 | 3.00E-01 | Haystead et al., 1992; Jain and Bhalla, 2014 |
| *ppMEK + ERK* $\boldsymbol{\leftrightarrow}$ *ppERK + ppMEK* | 3.24E-02 | 1.20E+00 | 3.00E-01 | Haystead et al., 1992; Jain and Bhalla, 2014 |
| *ppERK + MKP-1* $\boldsymbol{\leftrightarrow}$ *pERK+ MKP-1* | 1.50E-01 | 1.60E+01 | 4.00E+00 | Jain and Bhalla, 2014 |
| *pERK + MKP-1* $\boldsymbol{\leftrightarrow}$ *ERK+ MKP-1* | 1.50E-01 | 1.60E+01 | 4.00E+00 | Jain and Bhalla, 2014 |
