## Supplemental Table 4 for "Temporal pattern and synergy influence activity of ERK signaling pathways during L-LTP induction"

**Figure 1 –** **Source Data 4:** Reaction and rates constant involved in signaling pathways from calcium to CaMKII. Where indicated, CaMKII was optimized using De Koninck and Schulman, 1998 (Blackwell, 2019)

| Reaction equation | K_f_ (nM^-1^ Sec^-1^) | K_b_ (Sec^-1^) | K_cat_ (Sec^-1^) | Reference |
| --- | --- | --- | --- | --- |
| *Ca+ pmca* $\boldsymbol{\leftrightarrow}$ *pmca + Caext* | 5.00E-01 | 7.00E+00 | 3.50E+00 | Jȩdrzejewska-Szmek et al., 2017; Sedova and Blatter, 1999 |
| *Ca+ ncx* $\boldsymbol{\leftrightarrow}$ *ncx + Caext* | 1.68E-02 | 1.12E+01 | 5.60E+00 | Gall et al., 1999; Jȩdrzejewska-Szmek et al., 2017; Lőrincz et al., 2007 |
| *Caext +Leak* $\boldsymbol{\leftrightarrow}$ *Ca + Leak* | 1.50E-03 | 1.10E+00 | 1.10E+00 | Jȩdrzejewska-Szmek et al., 2017 |
| *Ca + Calbin* $\boldsymbol{\leftrightarrow}$ *CalbinCa* | 2.80E-02 | 1.96E+01 |  | Jȩdrzejewska-Szmek et al., 2017; Schmidt et al., 2007 |
| *CB + Ca* $\boldsymbol{\leftrightarrow}$ *CBCa* | 2.00E-02 | 1.00E+03 |  | Matthews et al., 2013; Matthews and Dietrich, 2015 |
| *Ng + CaM* $\boldsymbol{\leftrightarrow}$ *NgCaM* | 2.80E-02 | 3.60E+01 |  | Jȩdrzejewska-Szmek et al., 2017; Kubota et al., 2007 |
| *CaM+2Ca* $\boldsymbol{\leftrightarrow}$ *CaMCa2C* | 6.00E-03 | 9.10E+00 |  | Brown et al., 1997; Jȩdrzejewska-Szmek et al., 2017 |
| *CaMCa2C + 2Ca* $\boldsymbol{\leftrightarrow}$ *CaMCa4* | 1.00E-01 | 1.00E+03 |  | Jȩdrzejewska-Szmek et al., 2017; Putkey et al., 2003 |
| *CaM+2Ca* $\boldsymbol{\leftrightarrow}$ *CaMCa2N* | 1.00E-01 | 1.00E+03 |  | Brown et al., 1997; Jȩdrzejewska-Szmek et al., 2017 |
| *CaMCa2N + 2Ca* $\boldsymbol{\leftrightarrow}$ *CaMCa4* | 6.00E-03 | 9.10E+00 |  | Jȩdrzejewska-Szmek et al., 2017; Putkey et al., 2003 |
| *CaMCa4+CK* $\boldsymbol{\leftrightarrow}$ *CKCaMCa4* | 1.00E-02 | 1.50E+00 |  | Dupont and Goldbeter, 1998; Jȩdrzejewska-Szmek et al., 2017 |
| *2CKCaMCa* $\boldsymbol{\leftrightarrow}$ *CKpCaMCa4+ CKCaMCa4* | 3.83E-07 |  |  | Blackwell, 2019 |
| *3CKCaMCa* $\boldsymbol{\leftrightarrow}$ *CKpCaMCa4+ 2CKCaMCa4* | 3.56E-10 |  |  | Blackwell, 2019 |
| *4CKCaMCa* $\boldsymbol{\leftrightarrow}$ *CKpCaMCa4+ 3CKCaMCa4* | 2.24E-13 |  |  | Blackwell, 2019 |
| *2 CKpCaMCa4 + 2 CKCaMCa4* $\boldsymbol{\leftrightarrow}$ *3 CKpCaMCa4 + 1 CKCaMCa4* | 1.10E-15 |  |  | Blackwell, 2019 |
| *2 CKpCaMCa4 + 2 CKCaMCa4* $\boldsymbol{\leftrightarrow}$ *3 CKpCaMCa4 + 1 CKCaMCa4* | 3.03E-10 |  |  | Blackwell, 2019 |
| *2 CKpCaMCa4 + 2 CKCaMCa4* $\boldsymbol{\leftrightarrow}$ *3 CKpCaMCa4 + 1 CKCaMCa4* | 2.39E-10 |  |  | Blackwell, 2019 |
| *CKpCaMCa4* $\boldsymbol{\leftrightarrow}$ *CKp + CaMCa4* | 8.00E-04 | 1.00E-02 |  | Dupont and Goldbeter, 1998; Jȩdrzejewska-Szmek et al., 2017 |
| *CKp+PP1* $\boldsymbol{\leftrightarrow}$ *CK + PP1* | 4.00E-05 | 3.40E-01 | 8.60E-02 | Blackwell, 2019 |
| *CKpCaMCa4 + PP1* $\boldsymbol{\leftrightarrow}$ *CKCaMCa4 + PP1* | 4.00E-05 | 3.40E-01 | 8.60E-02 | Blackwell, 2019 |
| *Ip35 + PP1* $\boldsymbol{\leftrightarrow}$ *Ip35PP1* | 1.00E-03 | 1.10E-03 |  | Connor et al., 2000; Huang et al., 1999; Jȩdrzejewska-Szmek et al., 2017 |
