## Supplemental Table 5 for "Temporal pattern and synergy influence activity of ERK signaling pathways during L-LTP induction"

**Figure 1 –** **Source Data 5:** Reactions and rates constant involved in signaling pathways leading from cAMP to PKA

| Reaction equation | K_f_ (nM^-1^Sec^-1^) | K_b_ (Sec^-1^) | K_cat_ (Sec^-1^) | Reference |
| --- | --- | --- | --- | --- |
| *PKA + 2cAMP* $\boldsymbol{\leftrightarrow}$*PKAcAMP2* | 2.16E-04 | 6.00E-02 |  | Herberg et al., 1996; Jȩdrzejewska-Szmek et al., 2017; Ogreid and Døskeland, 1981 |
| *PKAcAMP2+2cAMP*$\boldsymbol{\leftrightarrow}$*PKAcAMP4* | 3.50E-04 | 6.00E-01 |  | Herberg et al., 1996; Jȩdrzejewska-Szmek et al., 2017; Ogreid and Døskeland, 1982 |
| *PKAcAMP4*$\boldsymbol{\leftrightarrow}$ *PKAr + PKAc* | 2.40E-01 | 2.55E-02 |  | Jȩdrzejewska-Szmek et al., 2017; Zawadzki and Taylor, 2004 |
| *PDE2+cAMP* $\boldsymbol{\leftrightarrow}$ *PDE2cAMP* | 2.00E-05 | 5.00E-01 |  | Blackwell et al., 2019 |
| *PDE2cAMP+cAMP* $\boldsymbol{\leftrightarrow}$*PDE2cAMP2 +AMP* | 5.90E-03 | 5.00E-01 | 5.40E+00 | Blackwell et al., 2019 |
| *PDE4+cAMP* $\boldsymbol{\leftrightarrow}$ *PDE4 +AMP* | 2.16E-02 | 6.90E+01 | 1.72E+01 | Herman et al., 2000; Jȩdrzejewska-Szmek et al., 2017 |
