## Supplemental Table 6 for "Temporal pattern and synergy influence activity of ERK signaling pathways during L-LTP induction"

**Figure 1 –** **Source Data 6:** Total concentrations of molecule species

| **Molecules** | **Concentration (nM)** |
| --- | --- |
| ERK | 360 |
| MEK | 680 |
| MKP1 | 40 |
| PP2A | 550 |
| BRaf | 300 |
| Raf1 | 300 |
| Rap1 | 400 |
| Ras | 600 |
| PDE2 | 160 |
| PDE4 | 100 |
| PKA | 1000 |
| Epac | 800 |
| Src | 20 |
| Cbl | 500 |
| CRKC3G | 500 |
| Sos | 100 |
| RasGRF | 400 |
| SynGap | 1000 |
| rasGap | 10 |
| rap1Gap | 12 |
| Cam | 28000 |
| CK | 20000 |
| PP1 | 3600 |
| Calbindin | 160000 |
| ncx | 2000 |
| pmca | 500 |
| CB (fixed buffer) | 3000000 |
| Ip35 | 500 |
| Ng | 21000 |
| Leak | 700 |
| Grb2 | 1000 |
